# *TOMM40* ‘523’ genotype induces sex- and tissue- specific differences in cholesterol and triglyceride levels in an *APOE-TOMM40* humanized mouse model

**DOI:** 10.64898/2026.06.09.731195

**Authors:** Neil V Yang, Dellila Hodgson, Timothy M Jang, Justin J Kim, William K Gottschalk, Hussein N Yassine, Ornit Chiba-Falek, Ronald M Krauss

## Abstract

**Introduction:** Genetic variants within the *APOE-TOMM40* locus are associated with Alzheimer’s disease (AD). A specific role for *TOMM40* is indicated by the finding that ‘523’ poly-T variants are associated with AD risk, but the mechanism for this effect has not been established. Our studies have shown that suppression of *Tomm40* in mice increased brain cholesterol content, an AD risk factor, and thus the present study sought to assess whether major ‘523’ poly-T variants (Short [S] and Very Long [VL]) are associated with altered lipid content of brain and other tissues.

**Methods:** We utilized a mouse model containing the entire human *APOE3-TOMM40* locus to quantify cholesterol and triglyceride levels in brain, liver, and white adipose tissue (WAT), as well as brain content of the AD biomarkers Aβ 42 and tau, in mice carrying two homozygous *TOMM40* ‘523’ poly-T genotypes (S/S and VL/VL).

**Results:** Male mice carrying the ‘523’-S/S genotype, but not females, showed higher brain cholesterol and triglyceride levels than VL/VL carriers, together with greater brain Aβ 42 content. WAT showed similar lipid differences as in the brain, while hepatic lipid content was broadly similar between ‘523’-S/S and -VL/VL genotypes, though there was a trend for higher triglycerides in VL/VL mice in a sex- and age-dependent manner.

**Discussion:** These results demonstrate that *TOMM40* ‘523’ poly-T variants drive tissue-specific, sex-, and age-dependent lipid differences in humanized *APOE3-TOMM40* mice, with the S/S genotype linked to elevated brain cholesterol and Aβ 42 levels, effects that link this locus to AD pathogenesis.

## INTRODUCTION

Alzheimer’s disease (AD) is the most common form of dementia, with 55 million people worldwide affected, and its prevalence is expected to triple to over 139 million by 2050^1^. In AD, factors including cholesterol imbalance and mitochondrial dysfunction promote amyloid beta (Aß) aggregation and phosphorylation of tau protein, two major hallmarks of AD pathology^2-5^. Thus, certain genetic variants of cholesterol and mitochondrial regulatory genes are associated with increased risk of late-onset AD (LOAD)^6-8^. Most notably, the APOE4 isoform of the gene encoding apolipoprotein E (APOE), a key lipid transport protein, is the strongest genetic risk factor for LOAD^9^. Despite the influence of the *APOE4* allele on AD risk, carriers of the *APOE3* allele are also at risk due to other genetic variants, including those within and neighboring the *APOE* locus on chromosome 19q13.32^10,11^.

Among these are polymorphisms within the translocase of the outer mitochondrial membrane 40 (*TOMM40*) gene which encodes the main channel-forming subunit of the translocase of the outer mitochondrial membrane (TOM) complex that is required for the transport of precursor proteins into the mitochondria to maintain mitochondrial function^12^. While most of the *TOMM40* polymorphisms are in linkage disequilibrium with variants in the *APOE* gene, the strong association with AD risk of a poly-T repeat within intron 6 (rs10524523, aka ‘523’) has been shown to remain significant after adjusting for the *APOE4* genotype^13-15^. This *TOMM40* ‘523’ variant has been classified into 3 distinct poly-T repeat lengths – short (S, T≤19), long (L, T=20-29), and very long (V≥, T30)^16^. Of these repeats, carriers of the ‘523’-L allele have been found to not only be significantly less prevalent than the S and VL alleles in the human population^13^, but also to be strongly linked to the *APOE4* isoform alone. The ‘523’-S and -VL alleles are typically linked with the *APOE3* allele^17,18^, and therefore determination of their associations with AD risk, independent of the influence of *APOE4*, is crucial. Notably, AD patients carrying genotypes derived from the ‘523’-S and -VL alleles have been reported to have differing associations with age at disease onset and cognition, such that ‘523’-S carriers experience faster cognitive decline than ‘523-VL carriers, and are therefore at a higher risk for AD than VL carriers^13,19,20^. While the basis for these relationships is not known, it has been shown that mice carrying the humanized ‘523’-S/S genotype have increased brain expression of human *APOE* mRNA vs those carrying the ‘523’-VL/VL allele^21^.

We have previously shown that suppression of *TOMM40* expression in human hepatocytes upregulates *APOE* gene expression due to activation of LXR (Liver X Receptors) by disruption of mitochondria-ER contact sites (MERCs), resulting in increased LDL receptor-mediated cholesterol uptake^22^. More recently we have found that LXR activation and MERC disruption are also induced by *TOMM40* KD in human-derived induced pluripotent stem cells (iPSCs) differentiated into neurons (iNeurons)^23^. Notably, *TOMM40* suppression in iNeurons promoted increased intracellular levels of cholesterol and Aß 42, markers of AD pathology, independent of the APOE isoform.

In the present study we utilized a mouse model in which the entire *Apoe-Tomm40* region was replaced with the human *APOE3* and adjacent *TOMM40* loci to assess the effects of the ‘523’ poly-T variants VL/VL and S/S on tissue lipid content, and to test whether these differ depending on age, sex, and tissue type.

## METHODS

### Humanized *APOE-TOMM40* mouse model

The mouse model used here was generated as described previously^21^ by replacement of the *Apoe-Tomm40* genomic region in C57BL/6J mice with the entire human *APOE3-TOMM40* locus, including all intronic and intergenic sequences and upstream and downstream regions. These mice contained *TOMM40* with either the homozygous “short (S)” allele at the ‘523’ poly-T site (T=15) or the homozygous ‘very long (VL)’ allele at this site (T=30). These genotypes were confirmed previously by DNA analysis^21^. To examine the influence of age on genotype effects, brain, liver, and white adipose tissue (WAT) samples were collected from mice of 6 and 12 months of age. The mice included in this study are presented in **Table S1**.

### Lipid extraction and quantification

Mouse brain, liver, and WAT samples were homogenized with a GentleMACS− dissociator (Miltenyi Biotec) in a chloroform:methanol:water solution (8:4:3, v/v/v) for total lipid extraction. Triglyceride (TAG) was quantified with the EnzyChrom− Triglyceride Assay Kit (BioAssay Systems) according to the manufacturer’s protocol. Lipid concentrations were measured at an absorbance of 570 nm. All samples were normalized to total protein concentration quantified by BSA assay (Genesee Scientific).

### Cholesterol extraction and quantification

Mouse brain, liver, and white adipose tissue (WAT) samples were homogenized with a GentleMACS− dissociator (Miltenyi Biotec) in a hexane:isopropanol (3:2, v/v) for cholesterol extraction. For cholesterol analysis, the samples were dried under nitrogen gas and then reconstituted in a buffer containing 0.5 M potassium phosphate (pH 7.4), 0.25 M NaCl, 25 mM cholic acid, and 0.5% Triton X-100. Intracellular cholesterol levels were subsequently measured using the Amplex Red Cholesterol Assay Kit (Life Technologies), following the manufacturer’s instructions. All samples were normalized to total protein concentration quantified by BSA assay (Genesee Scientific).

### Enzyme-linked immunosorbent assay (ELISA)

Isolated brain, liver, and WAT tissues were lysed in M Cellytic Lysis Buffer containing 1% protease inhibitor (Halt− Protease Inhibitor Cocktail; ThermoFisher Scientific) for 15 minutes using a cell disruptor or homogenizer. The lysates were then centrifuged at 14,000 × g for 15 minutes, and the supernatant was collected for further analysis. Aß 42 and total tau protein levels in the isolated tissues were measured using ELISA kits from ThermoFisher Scientific, following the manufacturer’s protocol. All results were normalized to the total protein content, determined by a BSA assay (Genesee Scientific).

### Statistical Analysis

Data are presented as mean ± standard error of the mean (SEM). The N-values in the figures represent biological replicates, with at least 5 replicates per group and experiment. For comparisons between two groups, statistical significance was determined using Student’s t-test. For comparisons involving more than two groups, one-way ANOVA or two-way ANOVA followed by Tukey’s post hoc test were used. All statistical analyses were performed with GraphPad Prism 9 (GraphPad Software, Inc.). A p-value of less than 0.05 was considered statistically significant.

## RESULTS

### *TOMM40* ‘523’ variant affects brain cholesterol dependent on poly-T length, sex, and age

We first measured cholesterol levels normalized to protein content in homogenized whole mouse brains and found no differences between the sexes or across age categories – 6 and 12 months (**Fig. 1A,B**). However mice with the ‘523’-S/S genotype had higher brain cholesterol levels than carriers of the VL/VL allele (**Fig. 1C**), though when stratified by sex, this was significant only in males (**Fig. 1D,E**). Further analysis showed that male mice carrying the *TOMM40* ‘523’-S/S allele had higher brain levels of free cholesterol and cholesterol ester vs VL/VL carriers only at 12-months of age (**Fig. 1F-K**). Thus, these findings show that the *TOMM40* ‘523’ genotype has effects on brain cholesterol levels in this mouse model that are dependent on sex and age.

**Figure 1.**
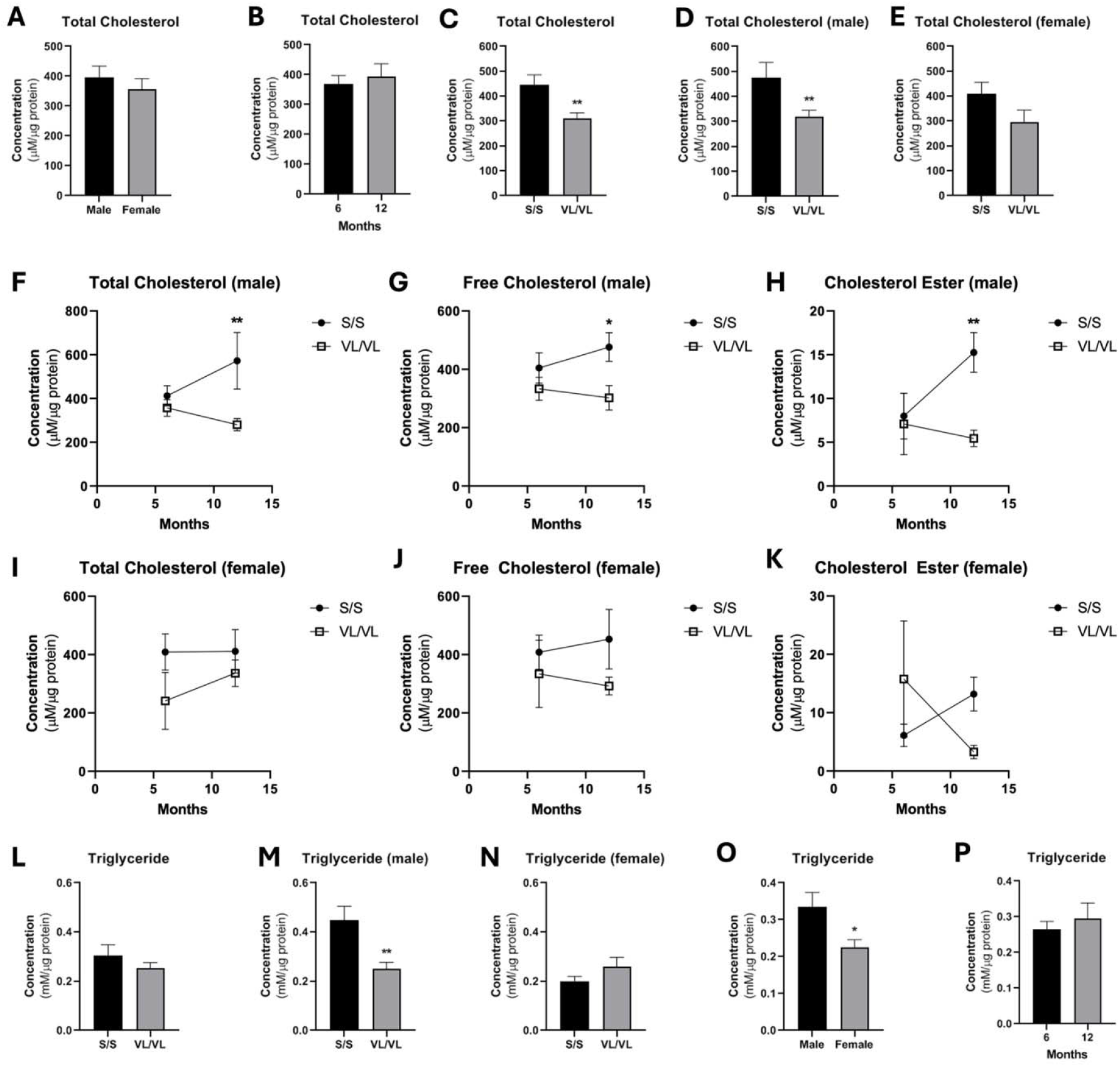
TOMM40 ‘523’-S/S genotype promotes brain cholesterol and triglyceride levels compared to *TOMM40* ‘523’-VL/VL genotype in male mice. Total cholesterol was extracted and quantified from whole brain homogenates of mice carrying the *TOMM40* ‘523’ -S/S vs VL/VL homozygous alleles, using the Amplex Red Cholesterol Assay: (A) Sex and (B) age did not affect total cholesterol levels. (C) Total cholesterol of S/S vs VL/VL in both sexes, (D) only males, and (E) only females were assessed. (F) Total cholesterol, (G) free, and (H) ester cholesterol levels in males were quantified across genotype and age. (I) Total cholesterol, (J) free, and (K) ester cholesterol levels in females were quantified across genotype and age. Triglyceride levels were quantified in whole brain homogenates of mice carrying the *TOMM40* ‘523’ -S/S vs VL/VL homozygous alleles: (L) Triglyceride of S/S vs VL/VL in both sexes, (M) only males, and (N) only females. (O) Sex affected brain triglyceride content, but not (P) age. For all: *n = 4-6* mice/group/sex. *\*p<0*.*05, **p<0*.*01* by one-way ANOVA, with post-hoc Student’s t-test to identify differences between groups. Data are represented as mean ± SEM.

Brain triglyceride content showed wide variation with no significant difference between the ‘523’-S/S and VL/VL genotypes in the combined sexes (**Fig. 1L**). However, similar to the cholesterol results, male mice with the ‘523’-S/S genotype had significantly higher brain triglyceride levels (**Fig. 1M**). While female mice showed the opposite trend, no significant differences were found without including age as a cofactor (**Fig. 1N**). Furthermore, female mice had lower brain triglyceride levels than males overall, but with no difference between 6 and 12 months of age (**Fig. 1O,P**). Together these results suggest that as for cholesterol, brain triglyceride content is influenced by *TOMM40* ‘523’ genotype dependent on sex and age.

We next assessed the effects of the *TOMM40* ‘523’ variants on brain content of Aβ 42 and total tau as biomarkers of AD pathogenesis and neuronal damage, respectively, in the context of human APOE3, independent of APOE4^24^. Male mice carrying the *TOMM40* ‘523’-S/S allele had higher brain Aβ 42 than those with ‘523’-VL/VL, similar to the results for cholesterol and triglyceride content (**Fig. 2A-E**). Males with ‘523’-S/S also had higher total tau levels than ‘523’-VL/VL though this did not reach statistical significance (**Fig. 2F-H**). Interestingly total tau levels decreased with age, with no changes between the sexes (**Fig. 2I,J**). These results indicate that in our humanized *APOE3-TOMM40* mouse model the effect of the *TOMM40* ‘523’-S/S variant on brain Aβ 42 content, a key marker of AD pathology, parallels the lipid effects of this variant, despite observing no differences in total tau levels in the brain.

**Figure 2.**
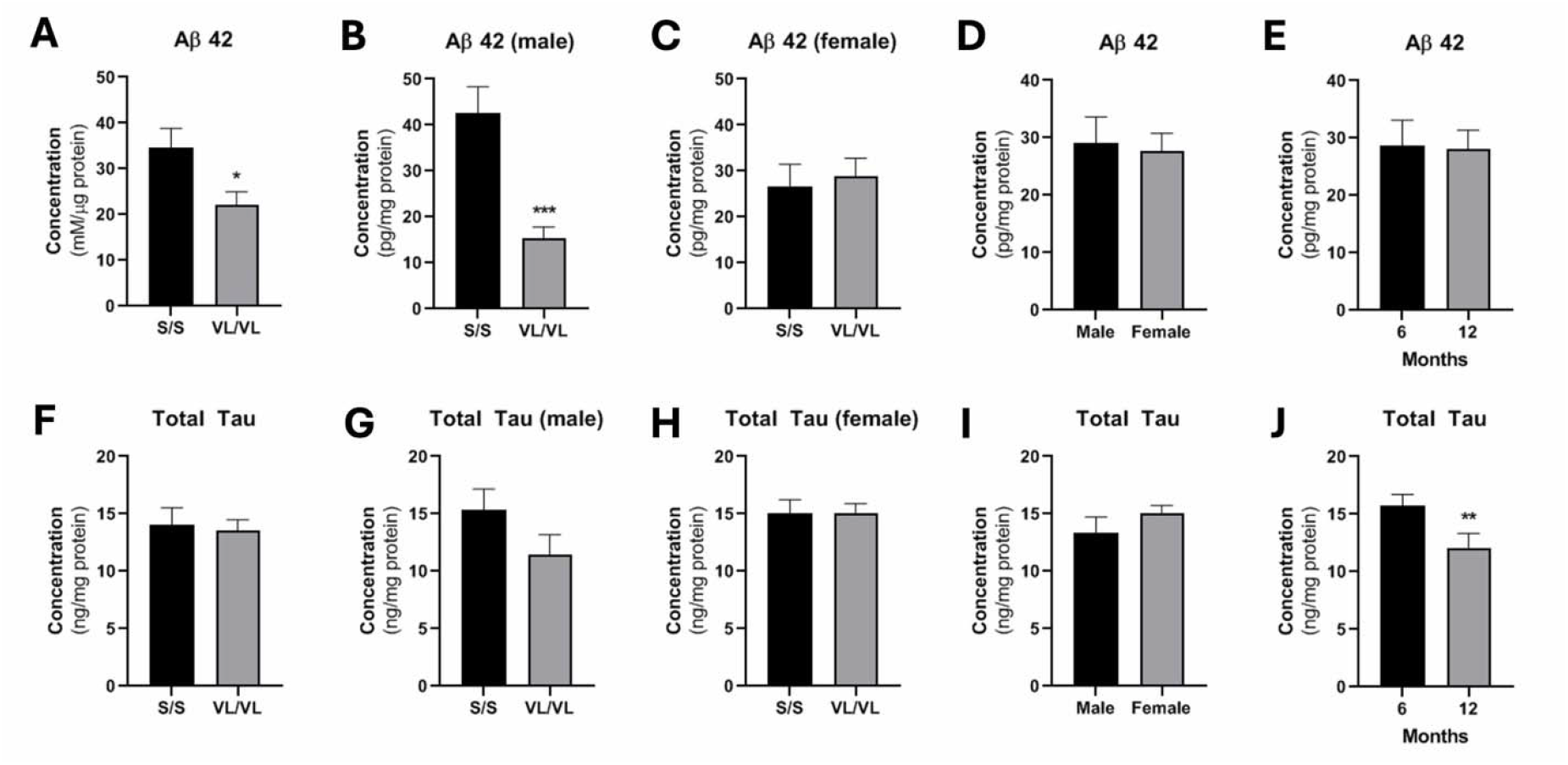
Aβ 42 levels in whole brain homogenates of male mice carrying the *TOMM40* ‘523’-S/S vs the ‘523’-VL/VL genotype. Aβ 42 levels in whole brain tissue were quantified in both sexes (A), male (B), and female (C) mice between *TOMM40* ‘523’ -S/S vs VL/VL genotypes. Sex (D) and age (E) did not affect total Aβ 42 levels. Total tau levels in whole brain tissues were quantified in the combined sexes (F), and separately in males (G), and females (H). There was no difference between the sexes (I), and in the combined group levels were lower at at 12 vs 6 months (J). For all: *n = 4-6* mice/group/sex. *\*p<0*.*05, **p<0*.*01, ***p<0*.*001* by one-way ANOVA, with post-hoc Student’s t-test to identify differences between groups. Data are represented as mean ± SEM.

### *TOMM40* ‘523’ poly-T genotype does not affect cholesterol levels in the liver

Consistent with previous studies, female mice had higher hepatic cholesterol content than males, possibly resulting from effects of estrogen on hepatic cholesterol synthesis, absorption, and/or degradation^22,25-27^ (**Fig. 3A**). In the combined sexes, liver cholesterol content was higher at 12 vs. 6 months of age, possibly due to increased cholesterol synthesis and/or uptake^28^ (**Fig. 3B**). There were no differences between the *TOMM40* ‘523’-S/S and’-VL/VL genotypes in either sex. (**Fig. 3C-E**). There were however transient spikes in cholesterol ester in the ‘523’-VL/VL males at 6-months of age (**Fig. 3F-H**) and in total and free cholesterol in females at 12-months (**Fig. 3I-K**). Hepatic triglyceride content was similar between the sexes (**Fig. 3L,M**). but interestingly it was substantially higher in mice with the ‘523’-VL/VL genotype though this did not reach statistical significance in females (**Fig. 3N-P)**.

**Figure 3.**
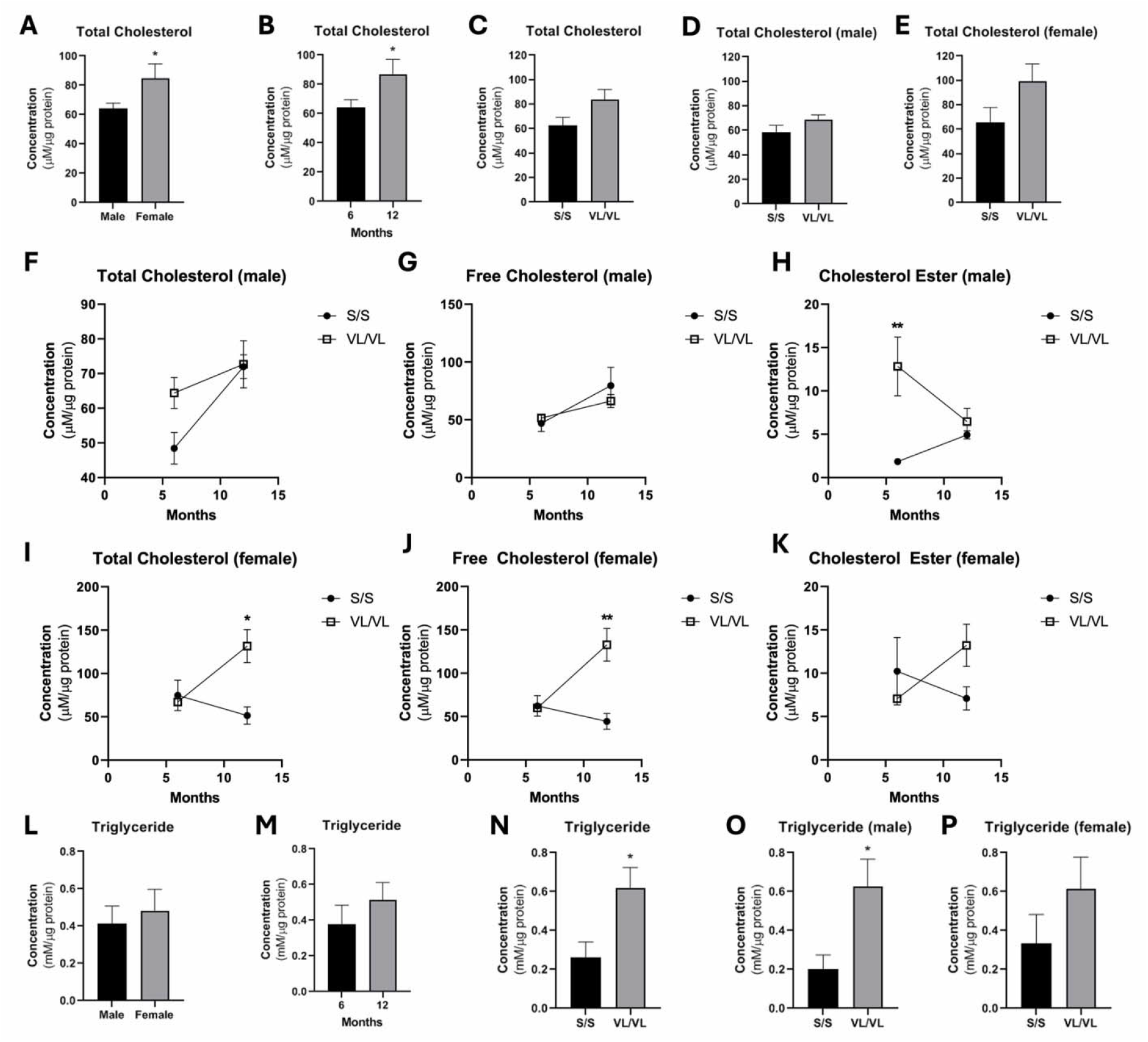
No difference in hepatic cholesterol but lower triglyceride content in male mice with the *TOMM40* ‘523’-S/S vs -VL/VL genotype. Total cholesterol and triglyceride were extracted and quantified from liver homogenates of mice carrying the *TOMM40* ‘523’ -S/S vs -VL/VL homozygous alleles, Total cholesterol levels stratified by (A) sex and (B) age. (C) Total cholesterol levels of S/S vs VL/VL in both sexes, (D) only males, and (E) only females. (F) Total cholesterol, (G) free, and (H) ester cholesterol levels in males across genotype and age. (I) Total cholesterol, (J) free, and (K) ester cholesterol levels in females across genotype and age. Sex (L) and age (M) did not affect liver triglyceride levels. (N) Triglyceride of S/S vs VL/VL genotypes in both sexes, (O) males, and (P) females. For all: *n = 4-6* mice/group/sex. *\*p<0*.*05* by one-way ANOVA, with post-hoc Student’s t-test to identify differences between groups. Data are represented as mean ± SEM.

### White adipose tissue cholesterol content is higher in mice with the *TOMM40* ‘523’-S/S vs. VL/VL genotype

Total cholesterol in WAT was similar between the sexes (**Fig. 4A**), with a non-significant trend toward lower levels with increasing age (**Fig. 4B**). As in the brain, total cholesterol was higher in *TOMM40* ‘523’-S/S mice compared to the ‘523’-VL/VL group in both males and females (**Fig. 4C-E**). Specifically, free cholesterol and cholesterol ester were higher at 12-months of age in male mice (**Fig. 4F-H**). Age-dependent increases in these measures were also seen in females (**Fig. 4I-K**). There were no differences in WAT triglyceride content between sex and age categories (**Fig. 4L,M**), but interestingly it was significantly higher in male *TOMM40* ‘523’-VL/VL carriers than ‘523’-S/S carriers, (**Fig. 4N-P**).

**Figure 4.**
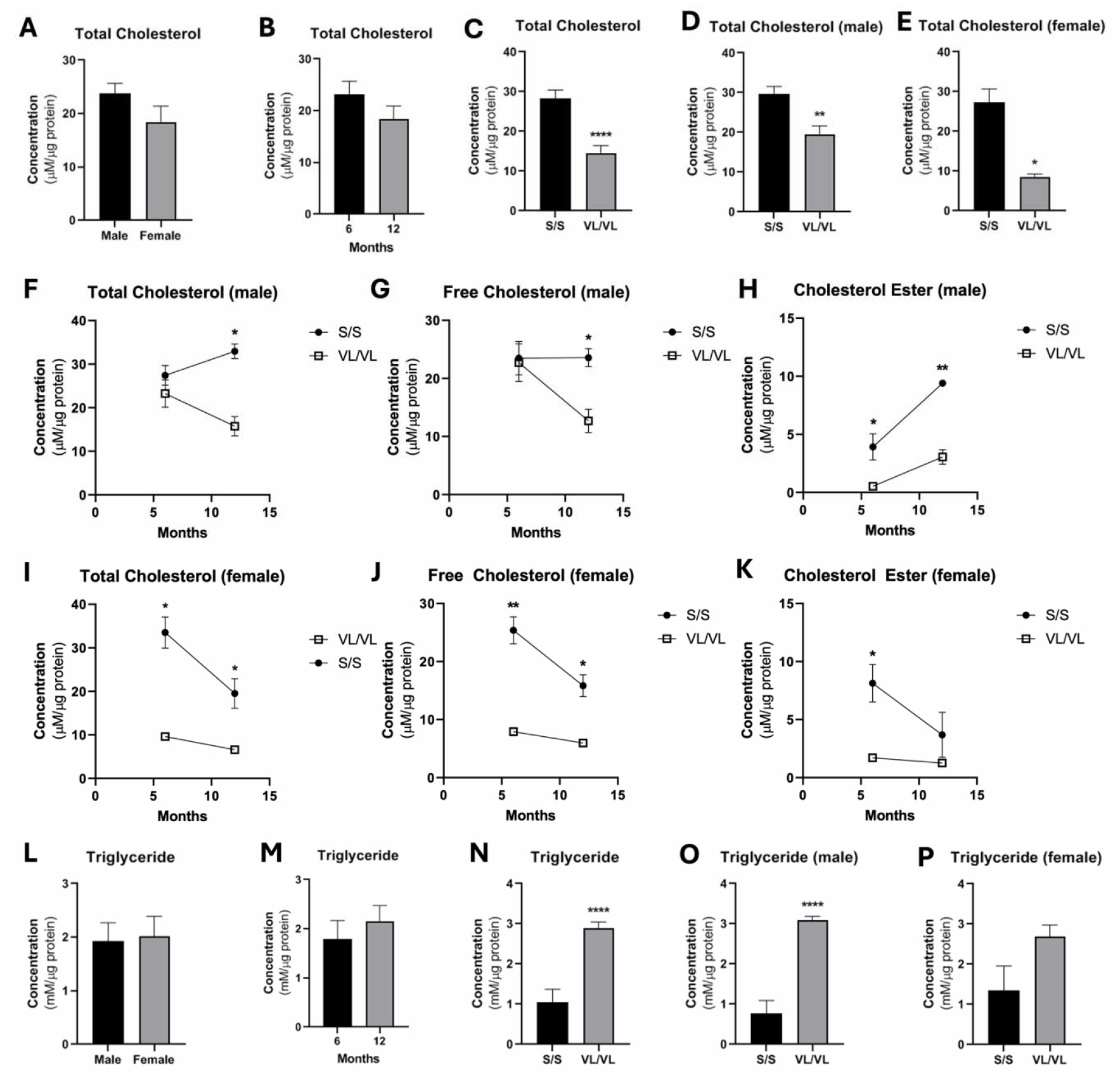
*TOMM40* ‘523’ genotype and sex influences cholesterol and triglyceride levels in white adipose tissue of *APOE3-TOMM40* ‘523’ mice. Total cholesterol was extracted and quantified from WAT homogenates of mice carrying the *TOMM40* ‘523’ -S/S vs -VL/VL alleles: (A) Sex and (B) age did not affect total cholesterol levels. (C) Total cholesterol of S/S vs VL/VL in both sexes, (D) males, and (E) females (F) Total cholesterol, (G) free, and (H) ester cholesterol levels in males across genotype and age. (I) Total cholesterol, (J) free, and (K) ester cholesterol levels in females across genotype and age. Sex (L) and age (M) did not affect WAT triglyceride levels. (N) Triglyceride content of WAT in S/S vs VL/VL genotypes of both sexes, (O) males, and (P) females. For all: *n = 4-6* mice/group/sex. \**\*p<0*.*01, ****p<0*.*00001* by one-way ANOVA, with post-hoc Student’s t-test was used to identify significant differences between groups. Data are represented as mean ± SEM.

## DISCUSSION

Previous GWAS studies in humans have shown that carriers of the *TOMM40* ‘523’-S/S genotype have accelerated development of AD pathology, including late-life cognition, compared to *TOMM40* ‘523’-VL/VL carriers^11,16,19,29^ and that its association with AD is not fully attributable to its linkage disequilibrium with APOE variants^30^. It has been reported that brain expression of *APOE* mRNA in a humanized mouse model is higher in mice with the *TOMM40* ‘523’-S/S vs. the VL/VL genotype^21^. The potential for this upregulation to impact brain cholesterol levels is indicated by our previous studies showing that KD of *Tomm 40* in mice results in increased APOE- and LDLR-mediated brain cholesterol content and that cholesterol levels in human iNeurons are increased by KD of *TOMM40* via this mechanism, independent of APOE isoform^23^. The present study aimed to test this possibility utilizing a humanized mouse model with the *TOMM40* ‘523’-S/S or ‘523’-VL/VL genotype encoded with the *APOE3/3* gene.

Our results confirm that brain cholesterol content is higher in mice carrying the human *TOMM40* ‘523’-S/S vs the VL/VL genotype, with the effect dependent on age and sex, such that it achieved significance only in males and was more pronounced with older age. Furthermore, we found higher brain levels of Aβ 42 in *TOMM40* ‘523’-S/S vs. VL/VL male mice. This may be attributed to a mechanism whereby increased saturation of cholesterol in the plasma membrane of neural cells enhances y-secretase cleavage of transmembrane amyloid precursor protein, leading to the production of Aβ 40/42 peptides, with Aβ 42 considered to be the primary form that drives neurodegeneration in AD^31-33^. Taken together with clinical evidence that increased brain cholesterol is a risk factor for AD^2,34^, the present findings suggest that the *TOMM40* ‘523’-S/S genotype may accelerate AD pathology through upregulation of brain cholesterol content.

We did not detect an effect of *TOMM40* ‘523’ genotype on brain content of total tau, which includes all isoforms of tau, including phosphorylated tau (p-tau). However, given that p-tau is the more accurate biomarker of AD risk, direct assessment of p-tau specifically in these mice will be required in future studies^35^. Unlike Aβ 42 aggregates, which appear early in AD pathogenesis, increased p-tau is detected in later stages^36-38^. Thus it would be informative in future studies to determine whether the *TOMM40* ‘523’-S/S genotype may be associated with increased brain p-tau, and in mice older than those studied here.

While women carrying *APOE-TOMM40* genetic variants have been reported to be at higher risk of developing AD than men^39^, effects of the *TOMM40* ‘523’-S/S genotype on AD biomarkers were significant only in our male humanized *APOE3-TOMM40* mice. This is consistent with our previous findings that KD of *Tomm40* in male mice increased brain cholesterol and Aβ 42 levels^23^, as well as hepatic lipid content and inducing hepatic steatosis^22^, while the opposite phenotypes and protective effects were observed in females. Particularly since pathways of lipid metabolism differ in mice vs. humans^40,41^ our findings point to the limitation of extrapolating the sexual dimorphism observed in our studies to humans, and the need to determine whether hormonal or genetic factors may be responsible.

Notably, we found that the differential associations of the *TOMM40* ‘523’ genotypes with brain cholesterol content were paralleled by those in WAT, but not in liver. Of potential relevance in this regard are our previous findings that although *TOMM40* KD promoted cholesterol uptake in both the brain^23^ and liver^22^, there was upregulation of genes promoting cholesterol efflux in liver which resulted in lowering of hepatic cholesterol content.

Our results showed that as for cholesterol, brain triglyceride content was higher in male mice with the *TOMM40* ‘523’-S/S vs. -VL/VL genotype, but this association was tissue specific in that triglyceride levels were reduced in WAT and liver. These results raise the possibility that in brain the *TOMM40* ‘523’-S/S genotype may operate to promote increases of cholesterol and triglyceride due to uptake of lipoprotein particles containing both lipids, though this mechanism and the tissue specificity and potential clinical significance of this observation will require further investigation.

A strength of the present study is the use of a humanized *APOE3-TOMM40* mouse model to identify associations of *TOMM40* ‘523’-S/S and VL/VL genotypes with lipid content of brain and other tissues. However, since our model was limited to assessing the *TOMM40* ‘523’-S/S and ‘523’-VL/VL genotypes, future studies will be needed to investigate the ‘523’ long/long (L/L) poly-T repeat genotype, which is exclusively linked to the *APOE4* isoform^42^. In addition, this study strongly supports our previous data^23^ whereby *TOMM40* ‘523’-S/S male mice with higher *APOE* mRNA expression levels exhibit upregulated cholesterol and Aβ 42 levels, compared to *TOMM40* ‘523’-VL/VL mice. This suggests that the differences in lipid and AD phenotypes between the two variants may be in part responsible for the early-onset and increased cognitive impairments in *TOMM40* ‘523’-S/S vs ‘523’-VL/VL carriers^13,19,20^, a possibility that will need to be further investigated. However, since *TOMM40* mRNA expression was previously shown to be higher in *TOMM40* ‘523’-S/S vs -VL/VL mice^21^, further study of this genetic variation’s effect on the transcriptional and translational regulation of *TOMM40/*TOMM40 will be required. Since genetic variants in poly-T tracts are known to affect mRNA splicing, thus impacting mRNA stability and translation efficiency^43,44^, it will also be necessary to determine whether the *TOMM40* ‘523’ genotypes influence the function and protein levels of TOMM40, and to identify downstream mechanisms by which such effects can impact tissue lipid levels.

In conclusion, this study demonstrates that *TOMM40* ‘523’ poly-T genetic variants are associated with differences in tissue lipid levels in a mouse model of the human *APOE-TOMM40* genetic locus, and that these differ by age, sex, and tissue type. The relevance of the results to AD pathogenesis is suggested by the finding of increased brain cholesterol and Aβ 42 levels in mice with the *TOMM40* ‘523’-S/S vs the ‘523’-VL/VL genotype, though the basis and pathophysiologic significance of the preferential effects in male mice will require further study.

## CONSENT STATEMENT

No human subjects were involved in this study.

## AUTHOR CONTRIBUTIONS

N.V.Y., O.C.F., and R.M.K. conceived the idea. D.H., W.K.G., and O.C.F. designed and created the humanized mouse model, performed the animal study and collected animal tissues. N.V.Y., T.J., and J.K. conducted the experiments and collected data. N.V.Y. analyzed the data and wrote the manuscript. D.H., H.Y., and O.C.F. assisted in data interpretation and editing of manuscript. R.M.K. carried out project supervision, data interpretation, manuscript writing and editing, and funding acquisition.

## DATA AVAILABILITY STATEMENT

Data will be provided upon reasonable request to the corresponding authors.

## SUPPLEMENTAL FIGURES

**Table S1.** Studied mice and tissue samples.

| **Tissue** | **Brain** |  |  |  |
| --- | --- | --- | --- | --- |
| ***TOMM40* '523' polyT genotype** | S/S |  | VL/VL |  |
| **Sex** | Male | Female | Male | Female |
| **Sample Size (n)** |  |  |  |  |
| 6 months | 6 | 6 | 6 | 6 |
| 12 months | 6 | 6 | 6 | 6 |
| **Tissue** | **White Adipose Tissue** |  |  |  |
| ***TOMM40* '523' polyT genotype** | S/S |  | VL/VL |  |
| **Sex** | Male | Female | Male | Female |
| **Sample Size (n)** |  |  |  |  |
| 6 months | 6 | 6 | 6 | 6 |
| 12 months | 6 | 6 | 6 | 6 |
| **Tissue** | **Liver** |  |  |  |
| ***TOMM40* '523' polyT genotype** | S/S |  | VL/VL |  |
| **Sex** | Male | Female | Male | Female |
| **Sample Size (n)** |  |  |  |  |
| 6 months | 6 | 6 | 6 | 6 |
| 12 months | 6 | 6 | 6 | 6 |

## REFERENCES

1. Reetz K, Liepelt-Scarfone I, Häger A, Flöel A, Schulz JB; Dementia Commission of the German Society for Neurology (DGN). The best treatment is prevention: prevention of cognitive decline and dementia - current state, gaps and next steps. Neurol Res Pract. 2026;8(1):30.

2. Chapleau M, Iaccarino L, Soleimani-Meigooni D, Rabinovici GD. The Role of Amyloid PET in Imaging Neurodegenerative Disorders: A Review. J Nucl Med. 2022;63(Suppl 1):13S–19S.

3. Ahmed H, Wang Y, Griffiths WJ, et al. Brain cholesterol and Alzheimer’s disease: challenges and opportunities in probe and drug development. Brain. 2024;147(5):1622–1635.

4. Wilkins HM. Interactions between amyloid, amyloid precursor protein, and mitochondria. Biochem Soc Trans. 2023;51(1):173–182.

5. Perez Ortiz JM, Swerdlow RH. Mitochondrial dysfunction in Alzheimer’s disease: Role in pathogenesis and novel therapeutic opportunities. Br J Pharmacol. 2019;176(18):3489–3507.

6. Li YJ, Nuytemans K, La JO, et al. Identification of novel genes for age-at-onset of Alzheimer’s disease by combining quantitative and survival trait analyses. Alzheimers Dement. 2023;19(7):3148–3157.

7. Grupe A, Abraham R, Li Y, et al. Evidence for novel susceptibility genes for late-onset Alzheimer’s disease from a genome-wide association study of putative functional variants. Hum Mol Genet. 2007;16(8):865–873.

8. Abraham R, Moskvina V, Sims R, et al. A genome-wide association study for late-onset Alzheimer’s disease using DNA pooling. BMC Med Genomics. 2008;1:44. Published 2008 Sep 29.

9. Fortea J, Pegueroles J, Alcolea D, et al. APOE4 homozygozity represents a distinct genetic form of Alzheimer’s disease. Nat Med. 2024;30(5):1284–1291.

10. Soyal SM, Kwik M, Kalev O, et al. A TOMM40/APOE allele encoding APOE-E3 predicts high likelihood of late-onset Alzheimer’s disease in autopsy cases. Mol Genet Genomic Med. 2020;8(8):e1317.

11. Xu J, Duan J, Cai Z, et al. TOMM40-APOE chimera linking Alzheimer’s highest risk genes: a new pathway for mitochondria regulation and APOE4 pathogenesis. Preprint. bioRxiv. 2024;2024.10.09.617477.

12. Gottschalk WK, Lutz MW, He YT, et al. The Broad Impact of TOM40 on Neurodegenerative Diseases in Aging. J Parkinsons Dis Alzheimers Dis. 2014;1(1):12.

13. Watts A, Haneline S, Welsh-Bohmer KA, et al. TOMM40 ‘523 Genotype Distinguishes Patterns of Cognitive Improvement for Executive Function in APOEL3 Homozygotes. J Alzheimers Dis. 2023;95(4):1697–1707.

14. Roses AD, Lutz MW, Amrine-Madsen H, et al. A TOMM40 variable-length polymorphism predicts the age of late-onset Alzheimer’s disease. Pharmacogenomics J. 2010;10(5):375–384.

15. Roses AD. An inherited variable poly-T repeat genotype in TOMM40 in Alzheimer disease. Arch Neurol. 2010;67(5):536–541.

16. Hayden KM, McEvoy JM, Linnertz C, et al. A homopolymer polymorphism in the TOMM40 gene contributes to cognitive performance in aging. Alzheimers Dement. 2012;8(5):381–388.

17. Watts A, Wilkins HM, Michaelis E, Swerdlow RH. TOMM40 ‘523 Associations with Baseline and Longitudinal Cognition in APOE L3 Homozygotes. J Alzheimers Dis. 2019;70(4):1059–1068.

18. Linnertz C, Anderson L, Gottschalk W, et al. The cis-regulatory effect of an Alzheimer’s disease-associated poly-T locus on expression of TOMM40 and apolipoprotein E genes. Alzheimers Dement. 2014;10(5):541–551.

19. Yu L, Lutz MW, Farfel JM, et al. Neuropathologic features of TOMM40 ‘523 variant on late-life cognitive decline. Alzheimers Dement. 2017;13(12):1380–1388.

20. Honea RA, Hunt S, Lepping RJ, et al. Alzheimer’s disease cortical morphological phenotypes are associated with TOMM40’523-APOE haplotypes. Neurobiol Aging. 2023;132:131–144.

21. Gottschalk WK, Mahon S, Hodgson D, et al. The APOE-TOMM40 Humanized Mouse Model: Characterization of Age, Sex, and PolyT Variant Effects on Gene Expression. J Alzheimers Dis. 2023;94(4):1563–1576.

22. Yang NV, Chao JY, Garton KA, et al. TOMM40 regulates hepatocellular and plasma lipid metabolism via an LXR-dependent pathway. Mol Metab. 2024;90:102056.

23. Yang NV, Wang S, Li B, et al. TOMM40 suppression promotes neuronal cholesterol imbalance and molecular and behavioral phenotypes of Alzheimer’s disease. Alzheimers Dement. 2026;22(4):e71306.

24. van Harten AC, Wiste HJ, Weigand SD, et al. Detection of Alzheimer’s disease amyloid beta 1-42, p-tau, and t-tau assays. Alzheimers Dement. 2022;18(4):635–644.

25. Conlon DM, Welty FK, Reyes-Soffer G, Amengual J. Sex-Specific Differences in Lipoprotein Production and Clearance. Arterioscler Thromb Vasc Biol. 2023;43(9):1617–1625.

26. Baptista CA, Didio LJ, Teofilovski-Parapid G. Variations of the blood supply of the human conus arteriosus. Bull Assoc Anat (Nancy). 1992;76(232):9–18.

27. Lorbek G, Perše M, Horvat S, Björkhem I, Rozman D. Sex differences in the hepatic cholesterol sensing mechanisms in mice. Molecules. 2013;18(9):11067–11085.

28. Nunes VS, da Silva Ferreira G, Quintão ECR. Cholesterol metabolism in aging simultaneously altered in liver and nervous system. Aging (Albany NY). 2022;14(3):1549–1561.

29. Chiba-Falek O, Gottschalk WK, Lutz MW. The effects of the TOMM40 poly-T alleles on Alzheimer’s disease phenotypes. Alzheimers Dement. 2018;14(5):692–698.

30. Yu L, Lutz MW, Wilson RS, et al. TOMM40’523 variant and cognitive decline in older persons with APOE ε3/3 genotype. Neurology. 2017;88(7):661–668.

31. Yang HS, Teng L, Kang D, et al. Cell-type-specific Alzheimer’s disease polygenic risk scores are associated with distinct disease processes in Alzheimer’s disease. Nat Commun. 2023;14(1):7659.

32. Banerjee S, Hashemi M, Zagorski K, Lyubchenko YL. Cholesterol in Membranes Facilitates Aggregation of Amyloid β Protein at Physiologically Relevant Concentrations. ACS Chem Neurosci. 2021;12(3):506–516.

33. Cruchaga C, Haller G, Chakraverty S, et al. Rare variants in APP, PSEN1 and PSEN2 increase risk for AD in late-onset Alzheimer’s disease families. PLoS One. 2012;7(2):e31039.

34. Ahmed H, Wang Y, Griffiths WJ, et al. Brain cholesterol and Alzheimer’s disease: challenges and opportunities in probe and drug development. Brain. 2024;147(5):1622–1635.

35. Hampel H, Blennow K, Shaw LM, Hoessler YC, Zetterberg H, Trojanowski JQ. Total and phosphorylated tau protein as biological markers of Alzheimer’s disease. Exp Gerontol. 2010;45(1):30–40.

36. Buchhave P, Minthon L, Zetterberg H, Wallin AK, Blennow K, Hansson O. Cerebrospinal fluid levels of β-amyloid 1-42, but not of tau, are fully changed already 5 to 10 years before the onset of Alzheimer dementia. Arch Gen Psychiatry. 2012;69(1):98–106.

37. Gibson Wood W, Eckert GP, Igbavboa U, Müller WE. Amyloid beta-protein interactions with membranes and cholesterol: causes or casualties of Alzheimer’s disease. Biochim Biophys Acta. 2003;1610(2):281–290.

38. Hernandez P, Lee G, Sjoberg M, Maccioni RB. Tau phosphorylation by cdk5 and Fyn in response to amyloid peptide Abeta (25-35): involvement of lipid rafts. J Alzheimers Dis. 2009;16(1):149–156.

39. Rajan KB, Weuve J, Barnes LL, McAninch EA, Wilson RS, Evans DA. Population estimate of people with clinical Alzheimer’s disease and mild cognitive impairment in the United States (2020-2060). Alzheimers Dement. 2021;17(12):1966–1975.

40. Zhu Q, Qi N, Shen L, et al. Sexual Dimorphism in Lipid Metabolism and Gut Microbiota in Mice Fed a High-Fat Diet. Nutrients. 2023;15(9):2175.

41. Gordon SM, Li H, Zhu X, Shah AS, Lu LJ, Davidson WS. A comparison of the mouse and human lipoproteome: suitability of the mouse model for studies of human lipoproteins. J Proteome Res. 2015;14(6):2686–2695.

42. Crenshaw DG, Gottschalk WK, Lutz MW, et al. Using genetics to enable studies on the prevention of Alzheimer’s disease. Clin Pharmacol Ther. 2013;93(2):177–185.

43. Hefferon TW, Broackes-Carter FC, Harris A, Cutting GR. Atypical 5’ splice sites cause CFTR exon 9 to be vulnerable to skipping. Am J Hum Genet. 2002;71(2):294–303.

44. Niksic M, Romano M, Buratti E, Pagani F, Baralle FE. Functional analysis of cis-acting elements regulating the alternative splicing of human CFTR exon 9. Hum Mol Genet. 1999;8(13):2339–2349.

